# Hypothalamic Farnesoid X Receptor deficiency alters energy balance by modulating hepatic glucose production and adipose tissue metabolism through central insulin signaling

**DOI:** 10.64898/2026.09.21.753141

**Authors:** Chloé Blondel, Cyril Bourouh, Emilie Nicolas, Aurélie Vadel, Emilie Dorchies, Emmanuelle Vallez, Anne Tailleux, Sophie Lestavel, Bart Staels, Kadiombo Bantubungi

**Author notes:** Corresponding authors (UMR1011, Laboratoire J&K, Boulevard du Pr. Jules Leclerc, 59045 Lille Cedex France) and (UMR1011, Institut Pasteur de Lille, 1 rue du Professeur Calmette, BP245, 59019 Lille France).

## Abstract

**Objectives:** The bile acid nuclear receptor Farnesoid X Receptor (FXR, NR1H4) is a major regulator of metabolism and energy homeostasis in peripheral organs. It modulates bile acid, glucose, and lipid metabolism, as well as fat mass and body weight. However, FXR is also expressed in the brain, particularly in the hypothalamus, a key center for the regulation of energy homeostasis. Although one study has demonstrated a role for brain FXR activation in energy balance, its specific hypothalamic role is still unknown. Here, we examined the role of FXR in the mediobasal hypothalamus in the regulation of energy balance.

**Methods:** We used a genetic approach combined with metabolic phenotyping to determine the effect of FXR invalidation in the mediobasal hypothalamus on metabolic parameters involved in the central regulation of energy homeostasis.

**Results:** Our results demonstrate that hypothalamic FXR deficiency induces a positive energy balance, resulting in a reduction in energy expenditure due to alterations in glucose metabolism accompanied by structural changes in white adipose tissues.

**Conclusion:** This study uncovers a previously unrecognized role for hypothalamic FXR in the central homeostatic control of energy balance, providing new insights into its contribution to peripheral glucose metabolism and adipose tissue structural remodeling.

## 1. INTRODUCTION

Energy homeostasis is a dynamic equilibrium between energy intake and energy expenditure, ensuring the maintenance of fat mass and body weight throughout life. Disruption of this balance leads to metabolic disorders such as obesity and type 2 diabetes. The central nervous system (CNS) plays a crucial role in regulating energy homeostasis, as hormonal and nutrient signals from peripheral organs converge on the CNS [1,2]. The CNS detects and integrates these signals to assess the body’s energy status and initiate appropriate biological responses. Among the key brain structures involved, the hypothalamus stands out as the primary center for the convergence and integration of peripheral signals, mainly due to two distinct neuronal populations in the arcuate nucleus: neurons co-expressing neuropeptide Y (NPY) and agouti-related protein (AgRP), and those expressing pro-opiomelanocortin (POMC) and cocaine- and amphetamine-related transcript (CART) [3].

Bile acids (BAs) are amphiphilic molecules synthesized from cholesterol in hepatocytes. BAs play a critical role in the intestinal absorption, digestion, and solubilization of fats, nutrients, and drugs. Beyond their digestive function, BAs also act as signaling molecules, particularly through their interaction with the Farnesoid-X-Receptor (FXR, NR1H4) and Takeda G protein-coupled receptor 5 (TGR5) [4,5]. FXR is a nuclear BA receptor primarily expressed in the liver and intestine but also found at lower concentrations in the white adipose tissues (WAT) [6]. Activation of FXR by BAs or synthetic ligands (such as GW4064 and tropifexor) leads to its binding to DNA response elements, thereby regulating various genes involved in bile acid, lipid, and glucose metabolism [7,8,9]. FXR also contributes to energy homeostasis; for instance, FXR knock-out mice (KO-FXR) exhibit reduced body weight gain and fat mass. In models of diet-induced or genetically modified obesity, FXR deficiency protects mice from excessive weight gain by decreasing fat mass in adipose tissue and improving glucose metabolism [10,11]. Conversely, peripheral administration of GW4064 enhances weight gain and glucose intolerance in diet-induced obesity [12]. These effects have been ascribed to FXR’s action on peripheral organs. Notably, FXR is also expressed in the brain, especially in the hypothalamus by the two above-mentioned neuronal populations of the arcuate nucleus, localized in the mediobasal hypothalamus (MBH) [13,14,15]. Deckmyn *et al*. initially demonstrated a role for brain FXR in regulating energy homeostasis by reducing energy expenditure through altered brown adipose tissue (BAT) function, mediated by changes in hypothalamic PKA-CREB signaling and sympathetic tone activity following intracerebroventricular injection of GW4064 [15].

However, the precise role of FXR in the hypothalamus has been overlooked. In the present study, the effect of hypothalamic FXR deletion on the regulation of energy homeostasis has been investigated using a genetic approach combined with metabolic phenotyping (cKO-FXR^MBH^). The findings reveal that cKO-FXR^MBH^ mice exhibited a positive energy balance accompanied by a reduction in hepatic glucose production during the pyruvate tolerance test, elevated lipogenesis and enhanced fatty acid uptake in both eWAT and scWAT.

## 2. MATERIALS AND METHODS

### 2.1. Animals and *in vivo* studies

The Institutional Committee approved animal experiments for the use and care of animals. The ethical committee at the University of Lille approved all protocols (APAFIS #45111- 2023061412504766 v11). Male FXR-deficient mice (*mus musculus*) (KO-FXR) (MGI:2137330) ; [16], male FXR-Floxed (FXR^fl/fl^) (MGI:6121372) and their littermate mice (10-12 weeks old) on the C57BL/6J background (Charles River), were housed under 12h light/12h dark cycles in temperature (21.5°C) and humidity-controlled rooms, in a specific pathogen-free environment. Mice were fed with a Standard diet (A04, Safe) with free access to water and food. Mice were placed in indirect calorimetry cages (TSE Systems, Hamburg, Germany) or standard cages, depending on the protocol.

#### 2.1.1. Whole-body FXR knockout mice (KO-FXR)

KO-FXR mice and their control (WT-FXR) were anesthetized using a mixture of ketamine (100mg/kg)/xylazine (20mg/kg) and stereotactically implanted with a cannula targeting the third ventricle of the brain (coordinates: +0,24mm rostral to bregma, +1mm lateral).. The cannula was secured on the skull with dental cement. Mice received an Intracerebroventricular (i.c.v) injection of insulin (0,005µg/µL, diluted in water) or vehicle 2 h prior to a pyruvate tolerance test (PTT). The solution (1 µL) was delivered at a rate of 0.5 µL/min. The PTT was performed after a 16 h fast by intraperitoneal injection of pyruvate (2 g/kg).

#### 2.1.2. Hypothalamic FXR deficient mice (cKO-FXR^MBH^)

FXR^fl/fl^ mice were randomized by body weight and age, then anesthetized (ketamine (100 mg/kg)/xylazine (20 mg/kg)). Bilateral stereotaxic injections of AAV vectors (pAAV.CMV.HI. eGFP-Cre.WPRE.SV40 (105545-AAV9, Addgene, RRID:Addgene_105545) or control pAAV.CMV.PI.EGFP.WPRE.bGH (105530-AAV9, Addgene, RRID:Addgene_105530)) were performed in the mediobasal hypothalamus (coordinates: −1.64 mm rostral to bregma, ±0.25 mm lateral, −5.7 mm depth). AAV vectors were diluted in PBS 0.001% pluronic acid. Each injection site received 0.5 µL containing 2 × 10^11^ vg/mL of each vector with a flow rate of 0.1 µL/min. The needle was left in place for 5 minutes to avoid vector reflux. At the end of the surgery, mice received a subcutaneous injection of anti-sedan. These mice were hereafter referred to as cKO-FXR^MBH^ and cWT-FXR^MBH^, respectively.

Glycemia was measured (AccuCheck Performa) in fed, fasting (2, 4, 6, and 24h), and refeeding (2, 4, and 6h) conditions. An intraperitoneal glucose tolerance test (IPGTT) was performed after a 4h fast by injection of glucose (1g/kg). Serum insulin levels were measured in basal condition and 15min after glucose administration using an ELISA kit (Mercodia, 10-247-01). A PTT was performed after a 16h fast by intraperitoneal injection of pyruvate (2g/kg). Triglycerides (TR210, Randox) and free fatty acids (157819910935, DiaSys) were measured using an INDIKO clinical chemistry analyzer (Thermo Fisher Scientific). To this end, mice were fasted for 5 h and then anesthetized with ketamine (100 mg/kg) and xylazine (20 mg/kg) before blood collection by retro-orbital puncture.

For energy metabolism analyses, cKO-FXR^MBH^ and their control cWT-FXR^MBH^ mice were individually placed in indirect calorimetry cages (TSE systems, Hamburg, Germany) for the measurement of energy parameters: food intake (grams), VO_2_ consumption (mL/h/kg lean mass), CO_2_ production (mL/h/kg lean mass), Respiratory Exchange Ratio (RER), locomotor activities (number of cages crossings) and energy expenditures (Weir formula: EE = (3.94xVO2 + 1.106xVCO_2_)/1000 (kcal/h/kg lean mass)).

### 2.2. Cell culture and treatments

Immortalized hypothalamic GT1-7 cells (Millipore Cat# SCC116, RRID:CVCL_0281) were cultured in DMEM GlutaMAX (61965-026, Gibco) supplemented with 10% fetal bovine serum (FBS; Gibco), 1 mM sodium pyruvate (11360-070, Gibco), 100 U/mL penicillin, and 10 mg/mL streptomycin (Gibco) at 37°C, 5% CO_2_.

FXR overexpression was achieved by electroporation using the Neon Transfection System (Life Technologies – 1350mV – 20ms), in the presence of 0.5 µg of empty plasmid or plasmid encoding FXRα1. The cell suspension was then transferred into antibiotic-free medium and seeded. GT1-7 cells were treated with insulin (200 nM, 10 min, diluted in water) or GW4064 (2µM, 4h, diluted in DMSO). Treatments were compared with vehicle controls (water or DMSO).

### 2.3. Western blot analysis

Cells were homogenized in 10µM Tris-320µMSucrose-1xPBS buffer and sonicated. Protein concentration was determined by the BCA-Protein Assay reagent kit. Protein in LDS sample buffer were loaded per lane, separated with NuPage 4-12% Bis-Tris Protein Gels (Thermofischer, NP0335BOX) and transferred to nitrocellulose membrane iBlot 2 transfert stacks (Thermofischer, IB23001). Membranes were immunoblotted at 4°C overnight with antibodies against Akt (Cell signaling, 9272S, RRID:AB_329827, rabbit polyclonal, IgG; 1/250), pAkt (Cell signaling, 9271S, RRID:AB_329825, rabbit polyclonal, IgG; 1/250) and GAPDH (SantaCruz, sc-25778, RRID:AB_10167668, rabbit polyclonal, IgG; 1/1000). All antibodies were diluted in Tris buffer saline (TBS) supplemented with Tween 0,05% and BSA 5%. The secondary antibodies used are anti-rabbit (Sigma A0545, RRID:AB_257896, IgG), diluted in TBS supplemented with Tween 0,05% and BSA 5% and incubated 2h at room temperature. Results are represented in the form of boxes for illustration purposes. All samples of an experiment were processed on the same western blot. When different gels were necessary, one or more common samples were run on each gel to allow subsequent normalization of the results.

### 2.4. Quantitative Real-Time PCR

Mediobasal hypothalamus, epididymal adipose tissue (eWAT), and subcutaneous adipose tissue (scWAT) of cKO-FXR^MBH^ and cWT-FXR^MBH^ were dissected and frozen in liquid N2. Total RNA was isolated using the RNeasy Lipid Tissue Mini Kit (Qiagen). Total RNA samples were extracted from GT1-7 cells by trizol reagent (Invitrogen). Retrotranscription reactions were performed using the cDNA Reverse Transcription High-Capacity Kit (Applied Biosystem). qPCR reactions were performed using Brilliant Sybr Green II QPCR Master Mix kit on the Applied Biosystems QuantStudio 3 (RRID:SCR_020238) (Supplemental Table 1). mRNA levels were normalized to a control gene (cyclophilin) whose expression is not influenced by the experimental conditions.

### 2.5. Immunohistochemistry

cKO-FXR^MBH^ and cWT-FXR^MBH^ mice were anesthetized with ketamine (100 mg/kg) and xylazine (20 mg/kg). Intracardiac perfusion was performed with 0.9% NaCl solution followed by 4% paraformaldehyde (PFA). Brains were post-fixed in 4% PFA for 4 hours at 4°C, then cryoprotected in successive 24-hour baths of sucrose-PBS buffer (10%, 20%, 30%) at 4°C. Brains were embedded in Frozen Section medium Neg-50, frozen in isopentane at -55°C, and stored at -80°C. Sections were cut at 40 µm using a Leica cryostat (CM3050) and stored in 12- well plates containing 3 mL PBS 0.01 M/azide 0.2%.

For free-floating immunofluorescence, brain sections of the mediobasal hypothalamus were placed in 12-well plates and incubated for 16 hours at 4°C with pAkt (Ser473) antibody diluted at 1/150 (3787, Cell signaling, RRID:AB_331170, rabbit monoclonal, IgG), CRE antibody diluted at 1/500 (MAB3120, Merck, RRID:AB_2085748, mouse monoclonal, IgG), GFAP antibody diluted at 1/500 (G3893, Sigma-Aldrich, RRID:AB_477010, mouse monoclonal, IgG), NEUN antibody diluted at 1/500 (ABN91, Merck Millipore, RRID:AB_11205760, chicken polyclonal, IgY) (PBS 0.01M + 0.1 or 0.3% Triton X-100 + 1% Normal Donkey Serum). Then, sections were washed three times with PBS 0.01M + 0.1 or 0.3% during 5 minutes and incubated for 2h at room temperature with the secondary antibody donkey anti-rabbit (for pAkt) (A10037, Thermofisher, RRID:AB_11180865) or donkey anti-mouse (for GFAP and CRE) (A10042, Thermofisher, RRID:AB_2534017) or donkey anti-chicken (for NeuN) (A78950, Thermofisher, RRID:AB_2921072) coupled to a fluorochrome emitting at 568 nm diluted at 1/200 (PBS 0.01M + 0.1% Triton X-100). Finally, labeling was performed using a 1/1000 diluted DAPI solution (62248, ThermoFisher) and slides mounted on Superfrost Ultra Plus Gold slides (11976299, Fisher Scientific) with Autofluorescent Eliminator Reagent (2160, Merck) and Dako® Fluorescent Mounting Medium (Agilent – S3023).

The image acquisition was performed using a confocal microscope (LSM 710 (RRID:SCR_018063) or LSM 930, Zeiss). The image analysis was performed using ImageJ software (RRID:SCR_003070).

### 2.6. Histology

cKO-FXR^MBH^ and cWT-FXR^MBH^ mice were anesthetized using a mixture of ketamine (100mg/kg)/xylazine (20mg/kg). Intracardiac perfusion of saline solution (NaCl 0.9%) was performed. eWAT, and scWAT were fixed for 24 hours in 4% PFA at 4°C, dehydrated, cleared, and embedded in paraffin. Paraffin blocks were cut at 5µm. Paraffin sections were stained with hematoxylin and eosin (H&E). The image acquisition was performed using a Zeiss Axioscan Z1 slide scanner (RRID:SCR_020927). Quantification was done using a macro in ImageJ software (RRID:SCR_003070) to calculate the number and size of adipocytes or lipid droplets.

### 2.7. Statistical analysis

Data were analyzed using the Mann-Whitney test, one-way ANOVA, and two-way ANOVA with Tukey’s *post hoc* using the Prism software (GraphPad, USA, RRID:SCR_002798)). All values are expressed as means ± SEM. Significance was set at p<0.05 for all experiments.

## 3. RESULTS

### 3.1. Development and validation of the hypothalamic FXR-deficient mice model

FXR invalidation was obtained by stereotaxic injection of FXR^fl/fl^ mice with a viral vector allowing CRE recombinase expression in the MBH (cKO-FXR^MBH^). Expression of GFP and CRE recombinase were analyzed by immunofluorescence in the arcuate nucleus of cKO-FXR^MBH^ and their controls (cWT-FXR^MBH^). As expected, CRE recombinase expression was observed solely in cKO-FXR^MBH^ mice (Figure Sup 1A, B). Next, *Fxr* expression was assessed by qPCR in the MBH and striatum (Figure Sup 1C, D). A significant decrease in *Fxr* mRNA level was found only in the MBH of cKO-FXR^MBH^ compared with their littermate controls, with no change in the striatum. Moreover, no difference in *Fxr* mRNA level in eWAT, and scWAT was observed, indicating an absence of peripheral spread of the AAV (Figures Sup 1E-G). Finally, using either NeuN or GFAP co-stainings, which are classical markers of neurons and astrocytes, respectively, the AAV-transduced GFP-positive cells were NeuN-positive and GFAP-negative, indicating the neuronal tropism of the used viral vector (Figure Sup 2A-D).

**Supplemental Figure 1:**
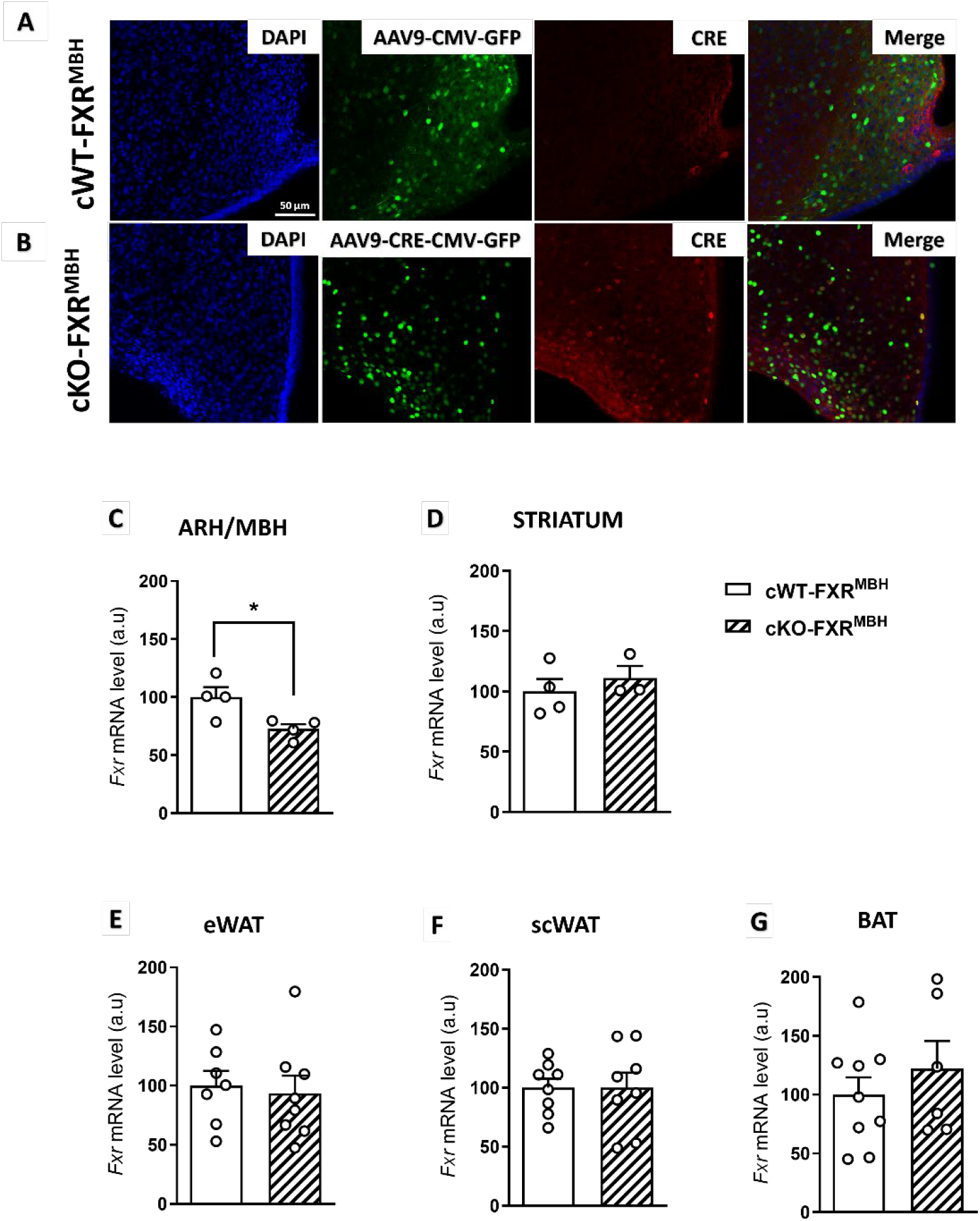
Targeted deletion of FXR in the mediobasal hypothalamus. (A-B) Representative images of CRE recombinase detected by immunofluorescence in the arcuate nucleus of cWT-FXRMBH (A) and cKO-FXRMBH (B) mice. The presence of endogenous fluorescence from the viral vector is shown in green, and DAPI counterstaining was performed (blue) (n=4/group). (C-G) Level of Fxr mRNA in the MBH (n=4/group) (C), the striatum (n=3-4/group) (D), the eWAT (n=7-8/group) (E), the scWAT (n=8/group) (F), the BAT (n=6-8/group) (G) of cKO-FXRMBH mice compared to cWT-FXRMBH mice. Data are expressed as mean ± SEM. Mann-Whitney test. *P < 0.05.

**Supplemental Figure 2:**
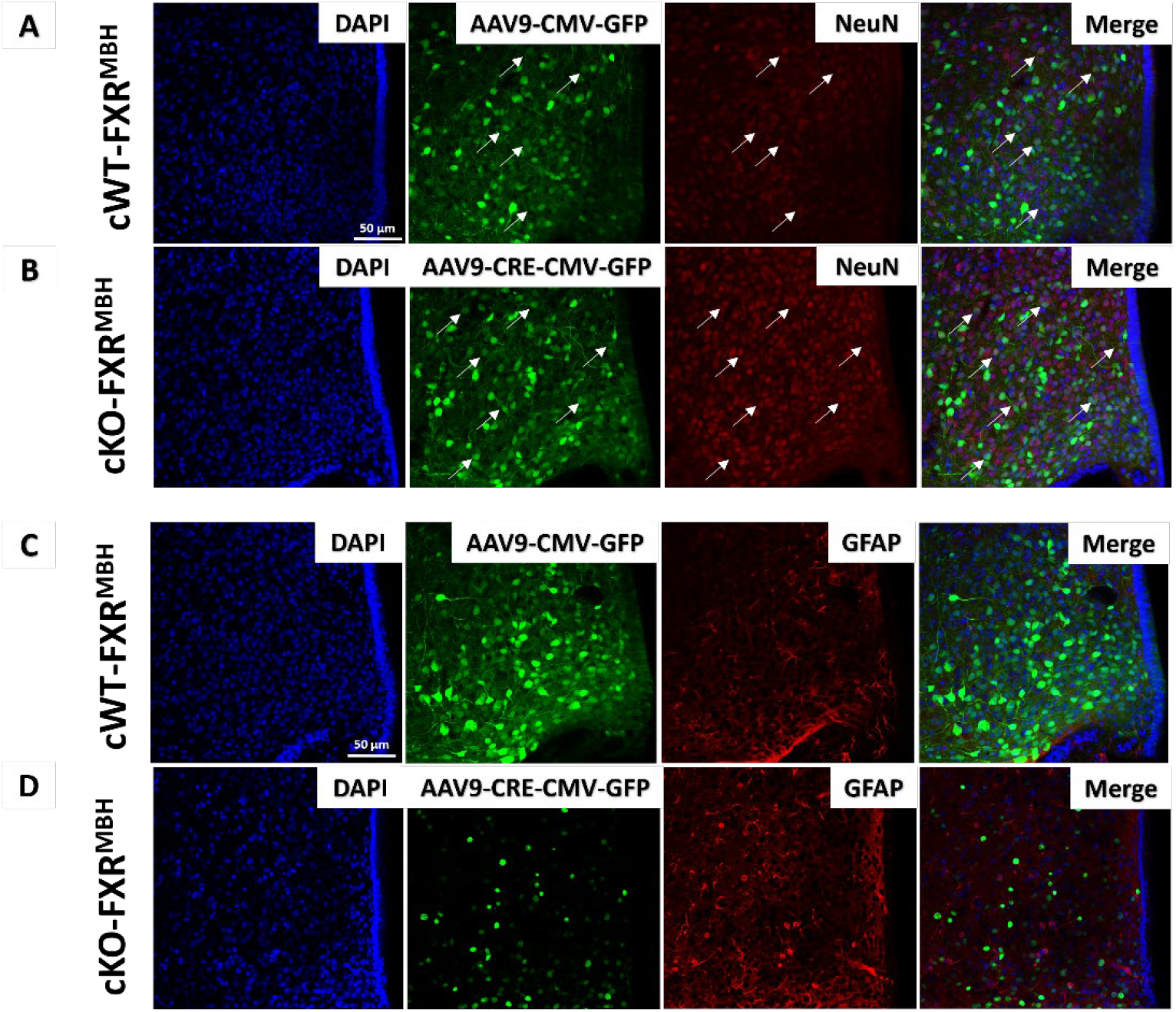
Neuronal tropism of AAV-mediated FXR deletion in the hypothalamus. (A-D) Representative images of NeuN (A-B) or GFAP (C-D) detected by immunofluorescence in the arcuate nucleus of cWT-FXR^MBH^ (A, C) and cKO-FXR^MBH^ (B, D) mice. The presence of endogenous fluorescence from the viral vector is shown in green, and DAPI counterstaining was performed (blue) (n=4/group).

### 3.2. Impact of hypothalamic FXR deficiency on energy homeostasis

To explore the impact of FXR knock-down in the MBH on energy homeostasis, body weights, food intake and energy expenditure were measured by indirect calorimetry in cKO-FXR^MBH^ and compared to cWT-FXR^MBH^ (Figure 1). Under *ad libitum* chow diet conditions, no difference in body weight was observed (Figure 1A). However, cKO-FXR^MBH^ mice showed increased food intake, reduced energy expenditure, and increased ambulatory activity as compared to cWT-FXR^MBH^ (Figure 1B-D). This latter observation likely explains the absence of body weight gain despite the increased food intake. Further, indirect calorimetry revealed a shift toward lipid utilization, as indicated by a decrease in VO_2_ and VCO_2_ and a lower respiratory exchange ratio (RER) in cKO-FXR^MBH^ mice as compared to their controls (Figure 1E-G). Taken together, these findings indicate that hypothalamic FXR knock-down disrupts energy homeostasis, leading to a positive energy balance despite unchanged body weight and increased ambulatory activity, which accounts for only a minor fraction (∼15%) of total energy expenditure [17].

**Figure 1:**
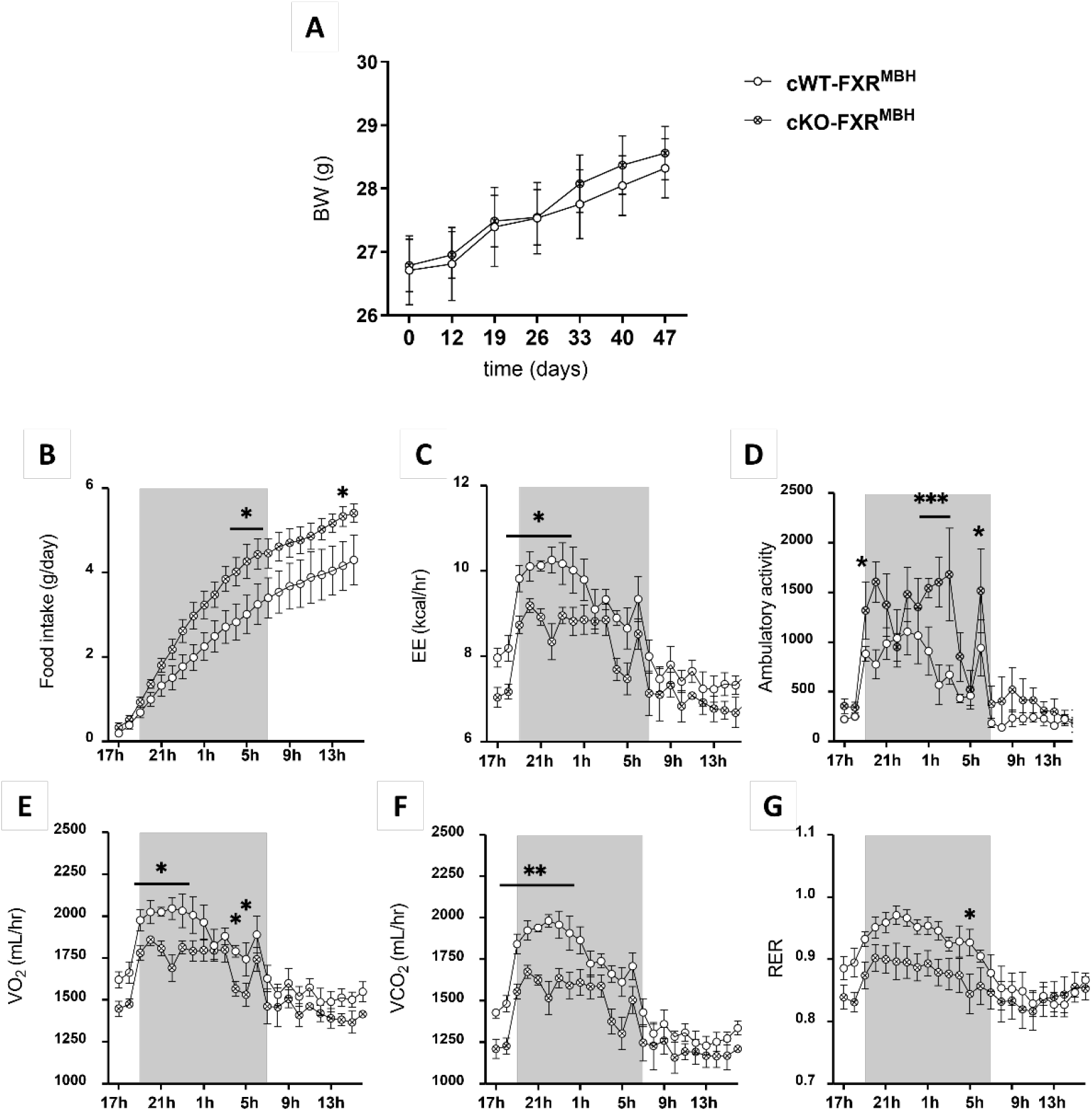
Hypothalamic FXR deficiency decreases energy expenditure. Body weight monitored every ten days for 50 days. (B-G) 24-hour indirect calorimetry analyses were performed in indirect calorimetry cages in cKO-FXR^MBH^ and cWT-FXR^MBH^, including food intake (B), energy expenditure (C), ambulatory activity (D), O_2_ consumption (E), CO_2_ production (F), and respiratory exchange ratio (RER) (G). (n=4/group). Data are expressed as mean ± SEM. *P < 0.05, ** P < 0.01, ***P < 0.001. Two-way ANOVA followed by Tukey’s post hoc test or Mann-Whitney test.

### 3.3. Role of hypothalamic FXR deficiency in glucose metabolism and lipid homeostasis

To explain the decrease in energy expenditure observed in cKO-FXR^MBH^ mice compared to control mice, we focused on basal metabolism, which accounts for 60% of energy expenditure [17]. First, we assessed glucose metabolism in cKO-FXR^MBH^ by measuring the dynamic glucose changes during fasting/refeeding, as well as using functional tests including an intraperitoneal glucose tolerance test (IPGTT) and a pyruvate tolerance test (PTT) (Figure 2). No difference between cKO-FXR^MBH^ and cWT-FXR^MBH^ in fasting and refeeding glycemia was observed (Figure 2A), nor in the IPGTT (Figure 2B-D). Moreover, cKO-FXR^MBH^ and cWT-FXR^MBH^ displayed a similar insulinemia in basal condition and 15 minutes after the initiation of the IPGTT test (Figure 2E). On the other hand, the PTT demonstrated a lower gluconeogenic potential in cKO-FXR^MBH^ mice compared to their controls (Figure 2F, G), associated with a tendancy of lower fasting plasma glucose level (Figure 2H).

**Figure 2:**
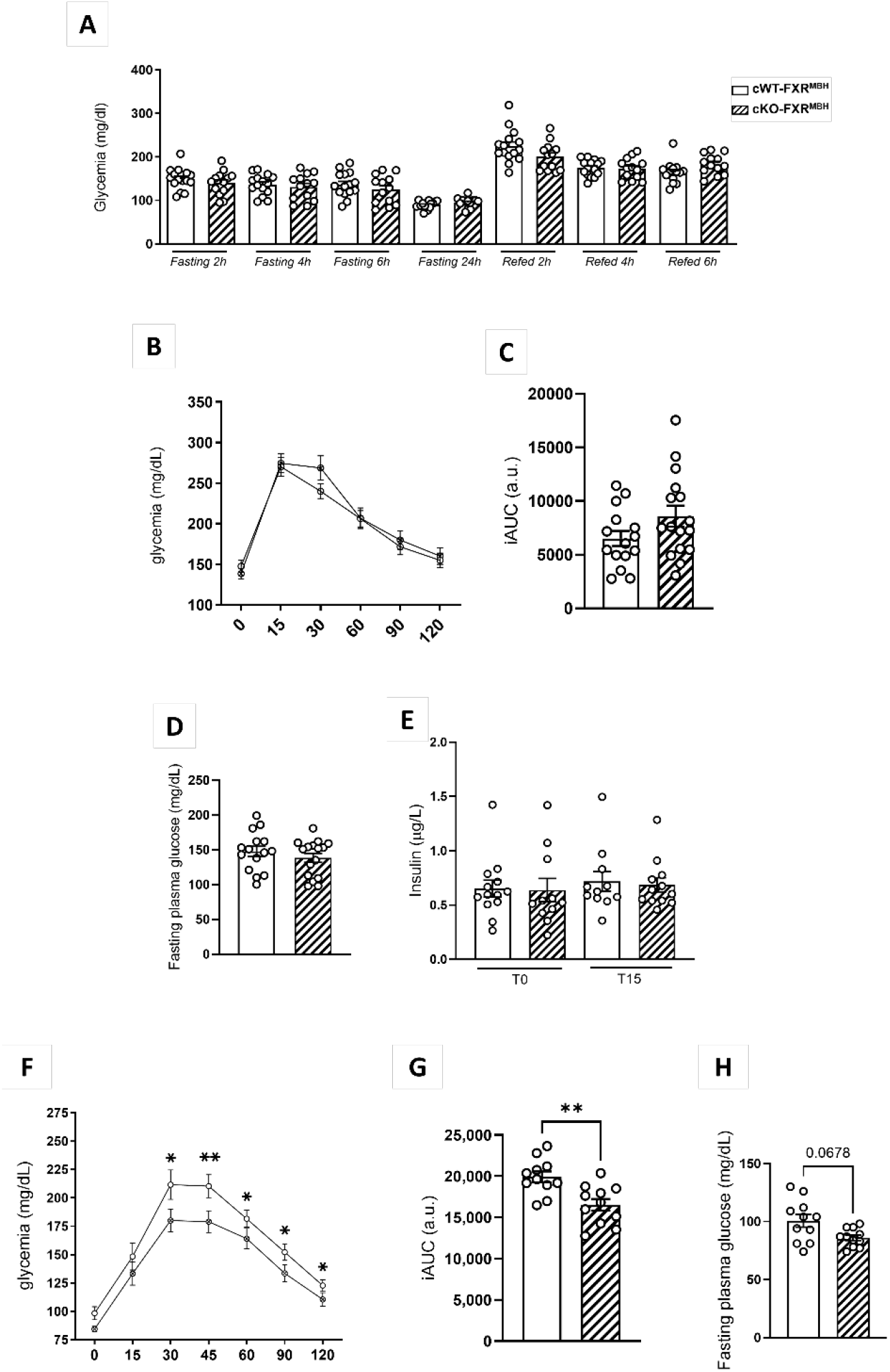
Hypothalamic FXR deficiency reduces hepatic glucose production but does not alter body weight gain, glycemia, or glucose tolerance. (A) Blood glucose monitoring at 2h, 4h, 6h, and 24h of fasting and at 2h, 4h, and 6h after refeeding (n=11-16/group). (B) Glucose excursion curve during IPGTT. (C) Calculation of the total iAUC of the IPGTT (n=15-16/group). (D) Fasting plasma glucose at the basal condition after the initiation of the IPGTT. (E) Serum insulin level at the basal condition (T0) and 15 min after the initiation of the IPGTT (T15). (F) Glucose excursion curve during PTT (n=11- 12/group). (G) Calculation of the total iAUC of the PTT. (H) Fasting plasma glucose at the basal condition after the initiation of the PTT. Data are expressed as mean ± SEM. *P < 0.05, **P<0.01. Two-way ANOVA followed by Tukey’s post hoc test or Mann-Whitney test.

Next, we examined the impact of hypothalamic FXR deficiency on lipid metabolism by studying WAT (Figure 3). While eWAT and scWAT mass was similar in both groups (Figure 3A, B, H, I), cKO-FXR^MBH^ mice displayed a significant increase in the lipid droplet size as compared to controls, in both eWAT and scWAT, as observed by histological and quantitative approaches (Figure 3C-F, J-M). Because such morphological changes suggest alterations in lipid metabolism, the gene expression of *de novo* lipogenic enzymes (stearoyl-CoA desaturase-1 (*Scd1*) and fatty acid synthase (*Fas*), diacylglycerol O-acyltransferase (*Dgat1)*), fatty acid uptake (*Cd36*) and lipolytic enzymes (hormone-sensitive lipase (*Hsl*), adipose triglyceride lipase (*Atgl*)) (Figure 3G,N) was evaluated. In eWAT, no changes in the expression of lipolytic enzymes (*Dgat1, Hsl, Atgl*) nor *Cd36* was observed between cKO-FXR^MBH^ and control mice. Conversely, an upregulation of *Scd1* and *Fas* in cKO-FXR^MBH^ (Figure 3G) was found, suggesting that hypothalamic FXR deficiency leads to increase *de novo* lipogenesis in eWAT. In scWAT, lipogenic gene expression was unchanged, while *Cd36* expression was significantly higher in cKO-FXR^MBH^ mice compared to control mice (Figure 3N). Moreover, plasma free fatty acids and triglycerides were lower in cKO-FXR^MBH^ mice as compared to cWT-FXR^MBH^ mice (Figure Sup 3A, B). Increased *Cd36* gene expression and these plasma changes suggest an enhanced fatty acid uptake and transport within scWAT adipocytes in cKO-FXR^MBH^ mice. Together, these findings, consistent with the observed decrease in energy expenditure, indicate that hypothalamic FXR deficiency influences both hepatic glucose production and adipose tissue lipid metabolism.

**Supplemental Figure 3:**
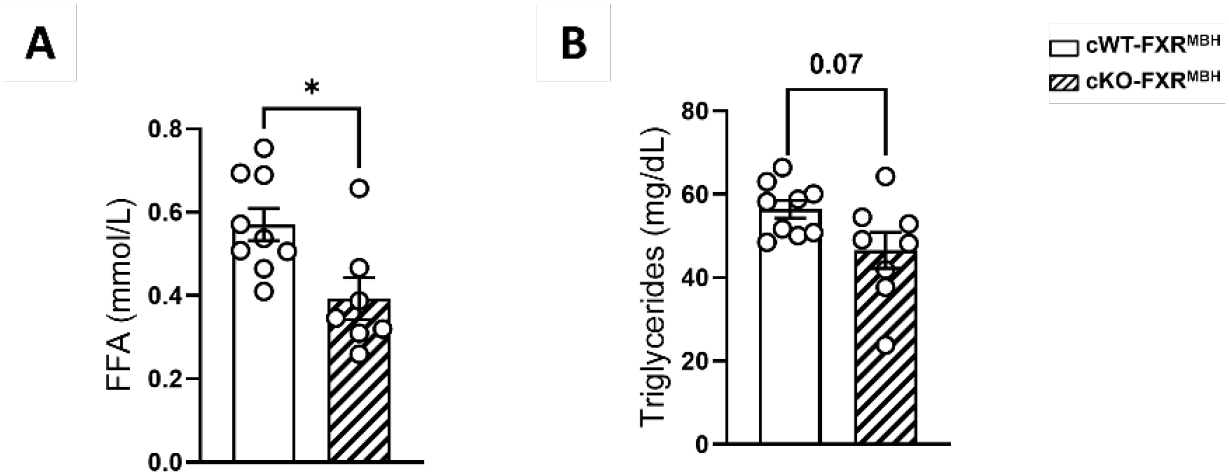
Plasma of FFA and TG in the hypothalamic FXR-deficient mice model. (A-B) Measurement of plasma free fatty acids (A) and triglycerides (B) after a 4h fast in cKO-FXR^MBH^ mice compared with cWT-FXR^MBH^ mice. Data are expressed as mean ± SEM. *P < 0.05. Mann-Whitney test.

**Figure 3:**
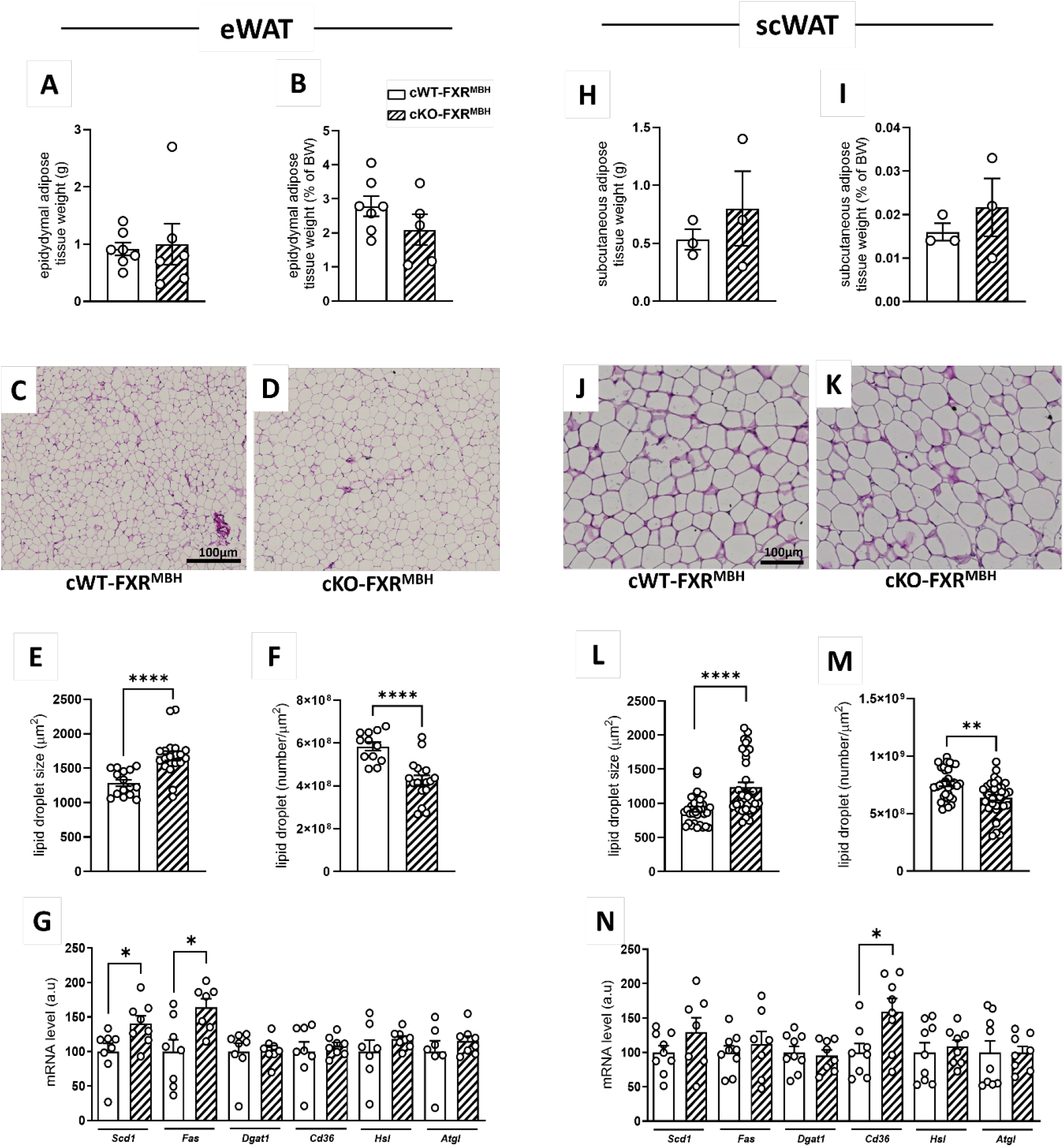
Impact of hypothalamic FXR deficiency on the structure and function of epididymal, subcutaneous and brown adipose tissues. (A) Weight of eWAT. (B) Weight of eWAT normalized to body weight. (C-D) Representative images of H&E histological staining of eWAT in cWT-FXR^MBH^ (C) and cKO-FXR^MBH^ (D) mice. (E-F) Quantification of the size (E) and number (F) of lipid droplets in eWAT. (G) mRNA Level of *Scd, Fas, Dgat1, Cd36, Hsl*, and *Atgl* by RT-qPCR in eWAT. (n=6-9/group). (H) Weight of scWAT. (I) Weight of scWAT normalized to body weight. (J-K) Representative images of H&E histological staining of scWAT in cWT-FXR^MBH^ (J) and cKO-FXR^MBH^ (K) mice. (L-M) Quantification of the size (L) and number (M) of lipid droplets in scWAT. (N) mRNA level of *Scd, Fas, Dgat1, Cd36, Hsl*, and *Atgl* by RT-qPCR in scWAT. (n=7-9/group). Data are expressed as mean ± SEM. *P < 0.05, ** P < 0.01, **** P < 0.0001. Mann-Whitney Test.

### 3.4. Role of hypothalamic FXR in brain insulin signaling

Notably, the energy and metabolic alterations observed in cKO-FXR^MBH^ are consistent with the well-established downstream effects of central insulin action [18]. We therefore hypothesized that FXR may modulate hypothalamic insulin signaling, which could then contribute to the observed metabolic phenotype. To investigate this hypothesis, we first assessed the impact of FXR overexpression (using an FXR expression plasmid delivered by electroporation; Figure 4A-C) or pharmacological activation (Figure 4D-F) on the insulin responsiveness of hypothalamic immortalized GT1-7 neurons. In response to insulin, the phosphorylation of pAkt was reduced upon both FXR overexpression or activation (Figure 4A-F), suggesting that FXR modulates the hypothalamic insulin response. Interestingly, it has been reported that brain insulin signaling suppresses neoglucogenesis [19,20]. In WT-FXR mice, i.c.v. insulin administration produced a modest reduction in glycemia during the test. In contrast, although KO-FXR mice displayed lower glycemia than WT-FXR mice under vehicle (VEH) conditions, i.c.v. insulin induced a greater suppression of glucose production in FXR-KO mice (Figure 4G- H), despite their lower baseline glycemia at T0 compared with control mice (Figure 4I). These findings indicate that FXR plays a role in modulating brain/hypothalamic insulin sensitivity, which is consistent with our *in vitro* observations. Finally, we evaluated the arcuate nucleus pAkt response to intraperitoneal insulin injection in whole body KO-FXR as well as in cKO-FXR^MBH^ mice and their controls. pAkt immunofluorescence upon insulin injection was higher within the arcuate nucleus in both whole body and cKO-FXR^MBH^ mice as compared to control mice (Figure 4J, K). These findings indicate that hypothalamic FXR plays a significant role in the response of the brain to insulin, particularly by enhancing insulin sensitivity.

**Figure 4:**
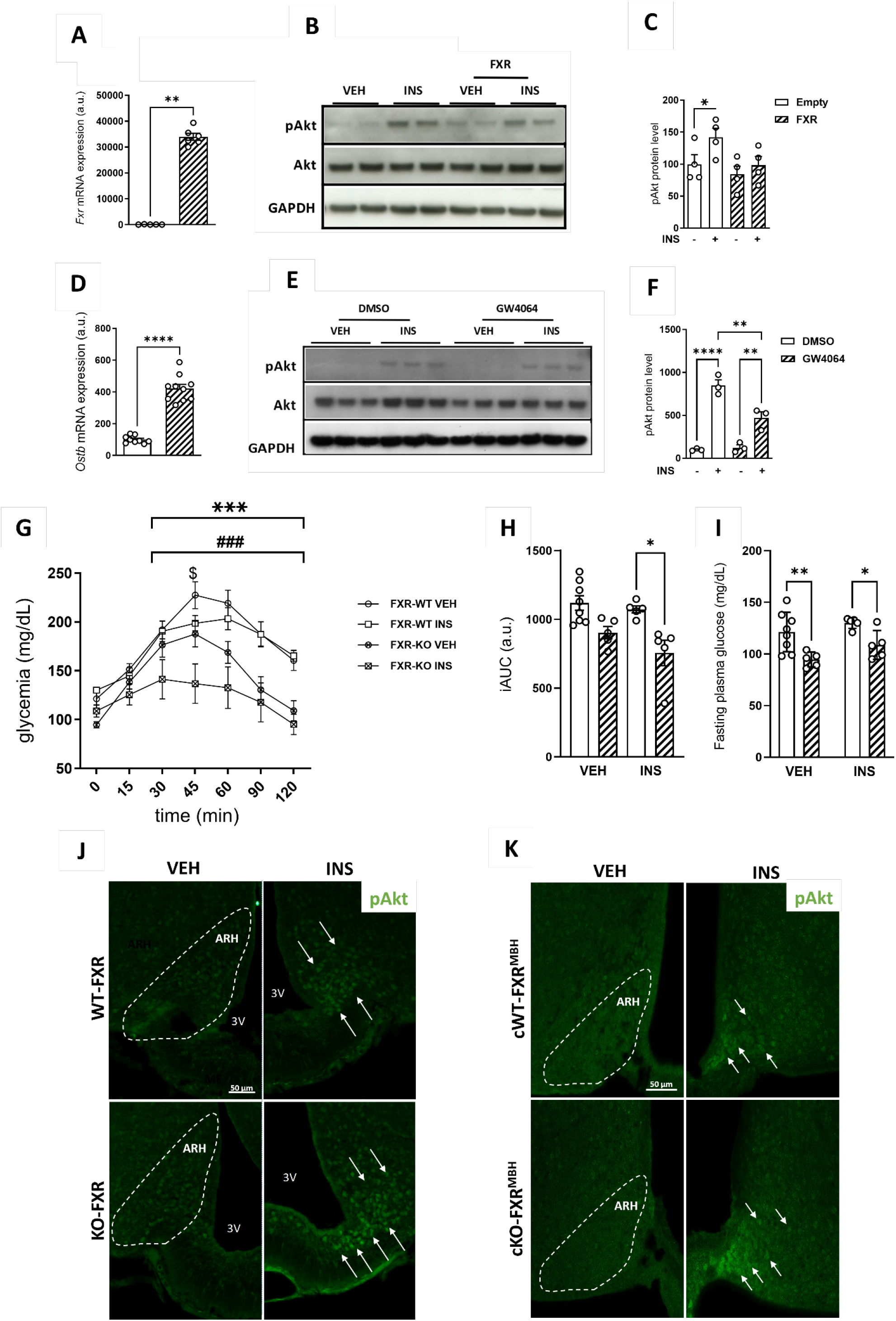
FXR deficiency improves central insulin response. *Fxr* mRNA level in GT1-7 following overexpression. (B) Protein expression of pAkt and Akt by western-blot following FXR overexpression in GT1-7. (C) Quantification of the pAkt/Akt ratio by Western blot. (D) mRNA level of *Ostβ*, a target gene of FXR, in GT1-7 following treatment with GW4064. (E) Protein expression of pAkt and Akt in GT1-7 treated with the agonist GW4064. (F) Quantification of the pAkt/Akt ratio by Western blot. (G) Pyruvate tolerance test performed on KO-FXR and WT-FXR mice following i.c.v injection of a vehicle solution (VEH) or insulin (INS). (H) Calculation of the total iAUC of the PTT. (I) Fasting plasma glucose at the basal condition after the initiation of the PTT. (J) Representative images of p(S473)Akt immunofluorescence in the arcuate nucleus following intraperitoneal administration of a vehicle solution (VEH) or insulin (INS) to KO-FXR and WT-FXR mice. (n=6/group). (K) Representative images of immunofluorescence of p(S473)Akt immunofluorescence in the arcuate nucleus following intraperitoneal administration of a vehicle solution (VEH) or insulin (INS) to cKO-FXR^MBH^ and cWT-FXR^MBH^ mice (n=4/group). Data are expressed as mean ± SEM. *P < 0.05, **P < 0.01, ***P < 0.001. Two-way ANOVA followed by Tukey’s *post-hoc* test or Mann-Whitney test. #: effect of insulin, *: effect of knock-out.

## 4. DISCUSSION

In the present study, we demonstrate that hypothalamic FXR contributes to the regulation of energy homeostasis, impacting feeding, energy expenditure, ambulatory activity and peripheral glucose and lipid metabolism. Furthermore, the concomitant decrease in VO_2_ and VCO_2_ observed through indirect calorimetry in mice indicates an overall reduction in oxidative metabolic rate. This systemic hypometabolism suggests reduced activity in all aerobic catabolic processes, including the oxidation of carbohydrates and lipids, thereby favoring the build-up of endogenous reserves by suppressing basal energy expenditure. In agreement with these observations, the effects of hypothalamic FXR deficiency led to decreased hepatic glucose production and increased endogenous lipid storage in WAT.

Indirect calorimetry analyses revealed that hypothalamic FXR deletion disrupts energy homeostasis. cKO-FXR^MBH^ mice displayed increased food intake and reduced energy expenditure, yet body weight remained unchanged. Since ambulatory activity contributes only modestly to total energy expenditure [17], factors beyond locomotor activity are likely contributing to the maintenance of body weight. Notably, cKO-FXR^MBH^ mice exhibited lower VO_2_, VCO_2_ and RER values during the dark phase, consistent with altered substrate utilization. Together, these findings indicate that hypothalamic FXR deletion induces a metabolic phenotype characterized by behavioral and metabolic traits commonly associated with increased susceptibility to obesity, despite the absence of overt body weight gain.

This study further identifies an impact of hypothalamic FXR deletion on eWAT and scWAT lipid storage regulation. While both tissues exhibit an increase in lipid droplet size, our data suggest that the mechanisms driving this change are distinct between the two tissues. Specifically, we observed an elevated expression of lipogenesis-related genes in eWAT, in favor of an expansion through synthesis, whereas the upregulation in scWAT may predominantly relate to fatty acid uptake, favoring expansion by influx. These results align with prior research [21-25], which demonstrates that WAT depots differ significantly in their structure, function, and metabolic profiles, largely due to the presence of unique adipocyte subpopulations. Taken together, our data suggest that hypothalamic FXR deficiency induces a pre-obese phenotype characterized by altered lipid handling despite unchanged body weight.

Collectively, our *in vitro* and *in vivo* data suggest that FXR overexpression or pharmacological activation impairs insulin signaling, whereas FXR deficiency enhances pathway activity. Given that insulin action on the brain favors lipid storage and suppresses gluconeogenesis [18-20], this perfectly aligns with the increased of lipid droplet size observed in adipose tissues, as well as the reduced fasting plasma glucose levels after overnight fasting and diminished gluconeogenic response during the pyruvate tolerance test in cKO-FXR^MBH^ mice. Together, these findings support an enhanced responsiveness to central insulin signaling following hypothalamic FXR deletion. Unexpectedly, while central insulin is typically considered an anorexigenic hormone, cKO-FXR^MBH^ mice also exhibit increased food intake. This surprising observation indicates that the full range of central insulin effects is not entirely reproduced following FXR suppression in the MBH. It is likely that certain insulin-mediated effects on food intake regulation depend on other hypothalamic or extra-hypothalamic neuronal subpopulations, as demonstrated by Bruning’s work [29-30].

Finally, hypothalamic BA-TGR5 signaling exerts an anorexigenic effect and protects against obesity [31-32]. Our findings support the notion that hypothalamic FXR knock-down triggers early metabolic alterations consistent with a pre-obese phenotype, potentially increasing susceptibility to obesity. In the hypothalamus, BA-FXR signaling might therefore exert similar outcomes as BA-TGR5 signaling. This is unexpected, given that both BA receptors often exert opposite actions in the peripheral tissues. To clarify this, a more precise measurement of BA in the hypothalamus could help reveal the underlying mechanisms. Further studies will be needed to better delineate the respective function of BA signaling in the hypothalamus and the brain.

In conclusion, our data demonstrate a novel role of FXR in the central regulation of energy homeostasis. These results pave the way for future studies on the function of hypothalamic FXR across different pathophysiological metabolic contexts. In addition, it seems also worthwhile to more generally evaluate the impact of BA changes in the hypothalamus in different metabolic contexts. Deepening our understanding of the neural circuits modulated by FXR and integrating these central actions with the receptor’s known peripheral effects, the potential to provide new effective therapeutic strategies to target FXR signaling in the treatment of metabolic disorders.

## CRediT authorship contribution statement

Chloé Blondel: Investigation, Formal analysis, Writing-original draft. Cyril Bourouh: Investigation. Emilie Nicolas: Investigation, Formal analysis. Aurélie Vadel: Investigation, Formal analysis. Emilie Dorchies: Investigation. Emmanuelle Vallez: Investigation, Formal analysis. Anne Tailleux: Resources. Sophie Lestavel: Resources. Bart Staels: Supervision, Funding acquisition. Kadiombo Bantubungi: Writing-original draft, Supervision, Methodology, Investigation, Formal analysis, Funding acquisition, Conceptualization.

## Acknowledgments

We thank Antonino Bongiovanni, Meryem Tardivel, Sarah Gabut, Brenda Lamens, and Solenne Audry of the BioImaging Center Lille (BiICEL) for their expertise and for training us on the used devices and the histological process. We thank the animal core facility of “Plateforme Lilloises en Biologie et Santé (PLBS) – UAR 2014 – US41” and Mélanie Besegher, A. Chatelain and Julien Devassine for mouse production and care. We thank Emilie Caron and Jessica Klucznik from Inserm UMR-S 1172 for their expertise on the use of indirect calorimetry cages. This work was supported by Inserm, ANR-17-CE14-0007 BABrain (B.S.), Labex EGID ANR10- LABX-46 (B.S.), Advanced ERC grant (to BS, Immunobile #694717) and EGID/I-SITE/Région Hauts de France and France Alzheimer.

